# Time course of EEG power during creative problem-solving with insight or remote thinking

**DOI:** 10.1101/2021.11.26.470102

**Authors:** Théophile Bieth, Marcela Ovando-Tellez, Alizée Lopez-Persem, Beatrice Garcin, Laurent Hugueville, Katia Lehongre, Richard Levy, Nathalie George, Emmanuelle Volle

## Abstract

Problem-solving often requires creativity and is critical in everyday life. However, the neurocognitive mechanisms underlying creative problem-solving remain poorly understood. Two mechanisms have been highlighted: forming new connections from and between the problem elements and *insight solving* (with a sudden realization of a solution). We examined EEG activity during an adapted version of a classical insight problem task, the Remote Associates Test, that requires finding a word connecting three words. It allowed us to explore *remoteness in semantic connections* (by varying the remoteness of the solution word across trials) and *insight solving* (identified as a “Eurêka” moment reported by the participants). *Semantic remoteness* was associated with a power increase in alpha band (8-12Hz) in a left parieto-temporal cluster, beta band (13-30Hz) in a right fronto-temporal cluster in the early phase of the task, and theta band (3-7Hz) in frontal cluster before the participants responded. *Insight solving* was associated with power increase preceding the response in alpha and gamma band (31-60Hz) in left temporal clusters and theta band in a frontal cluster. Source reconstructions show the brain regions associated with these clusters. Overall, our findings shed new light on the dynamic of some of the mechanisms involved in creative problem-solving.

## Introduction

Solving problems can be a societal challenge, an opportunity for progress, or a personal concern. We constantly have to find solutions to new problems and adapt ourselves to new situations, from the everyday life (e.g., how to reorganize my workspace at home), to worldwide concerns (e.g., how to avoid global warming). Problem-solving requires creativity (called here creative problem-solving) when there is no obvious or previously established rule to solve a newly encountered problem or when the heuristics or rules that we spontaneously use are inefficient or lead to an impasse. In creative problem-solving, we need to change our mental representation of the problem by recombining the elements of the problem in new ways or finding new connections between seemingly unrelated elements. In some cases, the solution comes to mind suddenly and spontaneously, with a “Eurêka” phenomenon (Topolinski & Reber, 2010). This problem-solving type is usually considered insight solving (Weisberg, 2013; Kounios & Beeman, 2014). It relates to the illumination phase of the creative process model developed from the reports of eminent scientific discoveries or artistic creations (Wallas, 1926). Combining remote elements and insight solving are considered as central aspects of creative thinking but the underlying neurocognitive mechanisms are still poorly understood. Are these two aspects related? What happens in the brain when solving a problem requires combining remote concepts or elicits a “Eurêka” experience? Here, we explore these questions using EEG during a problem-solving task assessing creative abilities.

Combining remote elements is a core component of the associative theory of creativity proposed by Mednick (Mednick, 1962). According to his approach, creativity relies on the ability to form new combinations from unusual associations. Mednick’s theory was operationalized in the Remote Associates Test (RAT) that consists in finding a word connecting three given unrelated cue words (Mednick, 1962). The RAT is a creative problem-solving task: it requires forming a new combination of distant elements of knowledge, and it often elicits an experience of insight or “Eurêka” in participants (Bowden et al., 2005; Topolinski & Reber, 2010; Kounios & Beeman, 2014). Several versions of the RAT have been developed using lexical (compound words) (Bowden & Jung-Beeman, 2003) or semantic associations between the cue words and the solution (Olteţeanu et al., 2019), or using pictures instead of words (Olteţeanu & Zunjani, 2020; Becker & Cabeza, 2021). Our lab developed a semantic associative version of the task (the Combined Associates Task, CAT) (Bendetowicz et al., 2017, 2018) in which we controlled the semantic association strength (*SAS*) between the expected solution and the three cue words. The CAT allows us to test Mednick’s hypothesis, according to which the more remote the elements to be combined, the more creative the process (Mednick, 1962).

A previous lesion study identified two distinct brain regions and networks as critical to CAT-solving when remoteness increases (Bendetowicz et al., 2018). First, the medial prefrontal cortex (PFC) as part of the default mode network, a network related to spontaneous cognition and associative thinking (Andrews-Hanna et al., 2010, 2014), was critical for the spontaneous generation of remote associates. Second, the rostro-lateral part of the PFC involved in the executive control network (Yeo et al., 2011; Power & Petersen, 2013) was critical for combining remote associates. These results are consistent with the associative theory of creativity but also emphasizes the importance of controlled processes during CAT-solving (Jones & Estes, 2015). They converge with findings from functional connectivity on divergent thinking in healthy subjects (Beaty et al., 2016), extend them to convergent thinking tasks (CAT), and demonstrate the necessity of both networks. Hence, their findings offer new light on the neural correlates of combining remote associates, while most previous neurocognitive studies that used RAT-like tasks focused on the insight phenomenon (Wu et al., 2020).

RAT-like tasks are helpful to explore insight solving because they provide multiple short trials, allowing to compare trials with and without insight, and better fit the constraints of neuroimaging studies than other insight problem-solving tasks (e.g., riddles). Currently, the subjective report of Eureka experience during problem-solving, on a trial-by-trial basis, is the most common measure used to study insight (Laukkonen & Tangen, 2018). The Eurêka corresponds to the subjective experience that arises when the solution comes to mind suddenly and effortlessly, without being able to report the mental steps leading to it. According to some insight theories (Sprugnoli et al., 2017), the Eurêka moment may follow an initial failure to solve the problem due to reaching a mental impasse and overcoming it with a reorganization of the problem representation (Ohlsson, 1992).

The critical question of the neural underpinnings of insight problem-solving remains unanswered. A few studies explored the brain correlates of insight problem-solving using functional MRI and reported the involvement of frontal regions (anterior and posterior cingulate cortex, inferior frontal gyrus), temporal regions (temporo-polar region, superior and middle temporal gyri, hippocampus) and the insula, during RAT-like tasks (Luo & Niki, 2003; Jung-Beeman et al., 2004; Anderson et al., 2009; Subramaniam et al., 2009; Aberg et al., 2016; Tik et al., 2018; Becker et al., 2020) or other insight tasks (Aziz-Zadeh et al., 2009; Dietrich & Kanso, 2010; Qiu et al., 2010; Shen et al., 2016; Lin et al., 2018). Electrophysiological methods such as EEG provide invaluable information on the time course of information processing and brain dynamics associated with cognitive processes. They thus have the potential to capture the suddenness of Eurêka experience (Jung-Beeman et al., 2004; Sandkühler & Bhattacharya, 2008). A pioneering study reported that RAT trials solved with Eurêka (compared to trials without Eurêka) were associated with a power increase in the alpha band in the right parieto-occipital areas around 1.5s before the subject’s response, followed by a gamma burst in the right antero-superior temporal lobe 0.3s before the subject’s response (Jung-Beeman et al., 2004). Alpha and gamma oscillations have been associated with insight solving in other studies that used the RAT (Sandkühler & Bhattacharya, 2008; Luft et al., 2018) and other paradigms (Sheth et al., 2009; Rosen & Reiner, 2016; Oh et al., 2020). Independently of insight solving, two studies reported a power increase in theta band in prefrontal electrodes and beta band in fronto-temporal electrodes when contrasting RAT-solving with a simple word generation task (Razumnikova, 2007) or a category fluency task (Danko et al., 2009).

Overall, the few existing neuroimaging studies of creative problem-solving focused mainly on insight, and none of them explored the effect of the remoteness of the elements to be combined. In addition, most EEG studies restricted their analyses to specific frequency bands or groups of electrodes. Hence, previous studies do not draw homogeneous conclusions on the brain mechanisms involved in creative problem-solving, including in RAT-like tasks. Here, we aim to better understand the neurocognitive mechanisms of creative problem-solving by jointly exploring the EEG correlates of the effects of *associative remoteness* and *insight solving*. For this purpose, we used the CAT (Bendetowicz et al., 2017, 2018), where the *remoteness* of the solution word varies across trials, and *insight* was explored by collecting subjective reports of Eurêka on a trial-by-trial basis. Since EEG data using the RAT are heterogeneous in the literature (Dietrich & Kanso, 2010) and the effect of *semantic remoteness* has not been investigated, we used an exploratory approach with no spatial, temporal, or frequency a priori. We hypothesized that the effects of *remoteness* and *insight solving* are associated with distinct brain EEG activities in space and time.

## Results

### Behavioral data

We recorded the EEG activity of 23 participants performing the CAT (100 trials). On each trial, participants had up to 30s to find a word that connects three unrelated words. Then they reported if they solved the trial with a Eurêka (**Figure 1**; see *method*). Each trial was characterized by a semantic association strength (*SAS*) value (a continuous variable determined by the material and fixed between subjects) and categorized according to how the subject solved it (with or without Eurêka; binary variable that depends on each subject).

**Figure 1.**
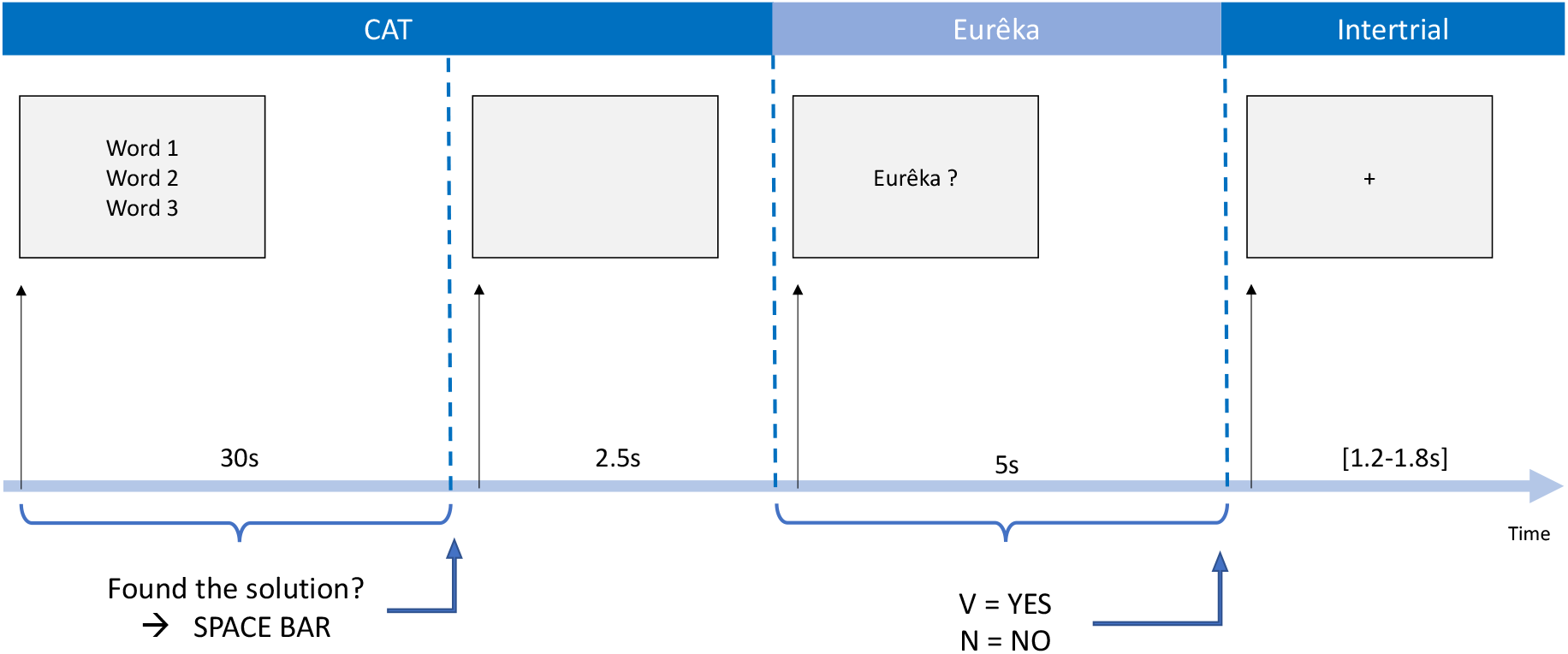
Summary of the CAT procedure. Experimental design of the CAT. Each trial starts with the presentation of three unrelated words, vertically displayed on a grey screen for up to 30s. The participants press the space bar as soon as they think they have the solution, triggering the display of a blank screen during 2.5s. They verbalize their response during this period. Then, the question “Eurêka?” is displayed on the screen, and the participants indicate whether the solution that they just gave came to their mind with a Eurêka, using the keyboard letters “V” (yes) and “N” (no), within a time limit of 5s. Finally, a fixation cross is displayed on the screen for a random time before beginning a new trial (intertrial interval ranges between 1.2 and 1.8s).

Overall, mean accuracy across individuals was 57.4% (SD=12.0), and mean RT was 8.4s (SD=1.0).

Across trials, the percentage of participants who gave a correct response correlated significantly positively with *SAS* (**Figure 2A**, rho=0.48, *p*=3.85 10^−7^), indicating that the closer the solution was, the more individuals found it. The correlation between the mean RT for correct responses across trial and *SAS* was negative and marginally significant (**Figure 2B**, rho=-0.20, *p*=0.051).

**Figure 2.**
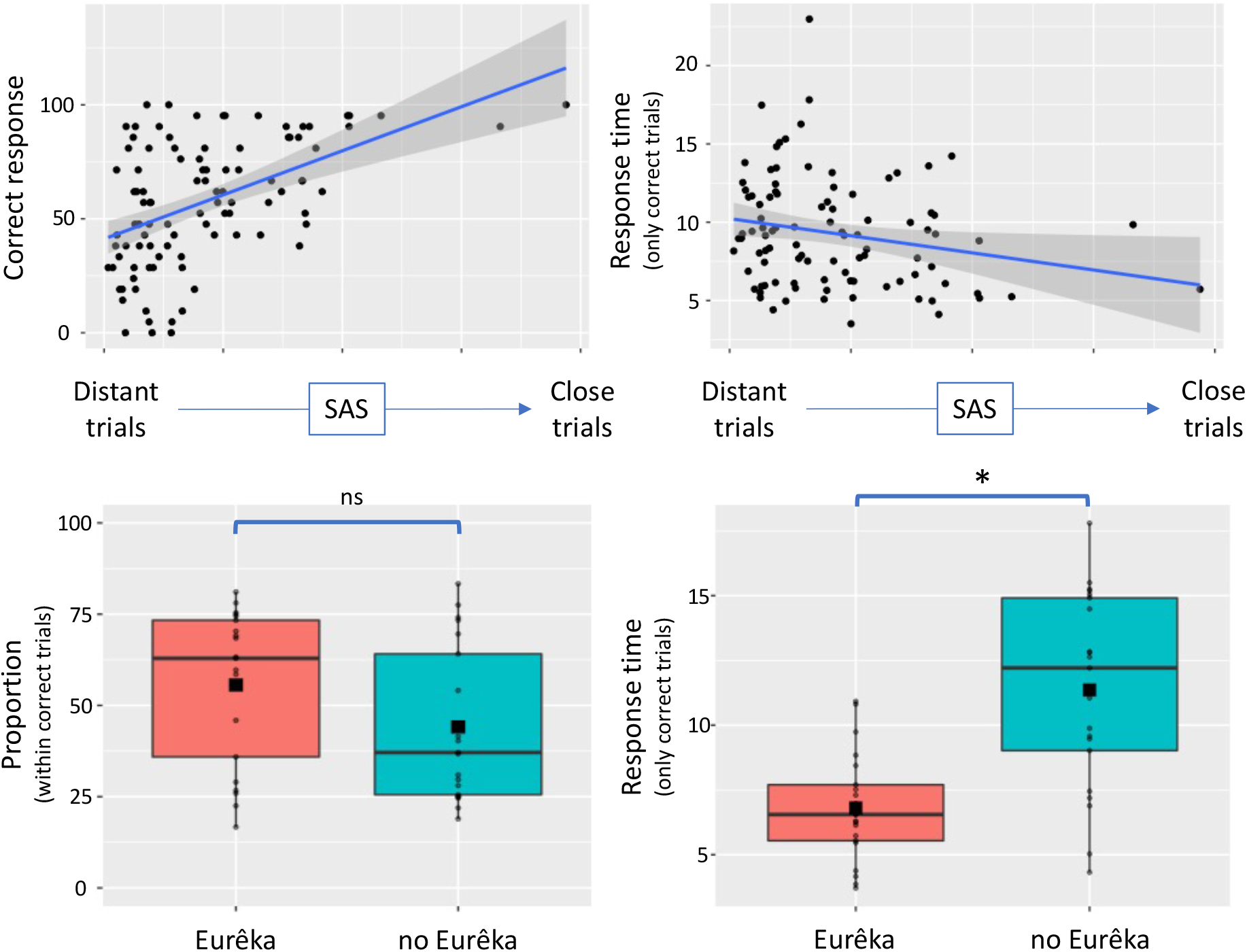
Behavioral results. **A**. Percentage of participants with correct responses per trial as a function of *SAS*. Each dot represents a trial, and the blue line represents the regression line between the two variables (rho=0.48, *p*=3.85 10^−7^). **B**. Averaged RT per trial as a function of *SAS*. Each dot represents a trial, and the blue line represents the regression line between the two variables (rho=-0.20, *p*=0.051). **C**. Percentage of Eurêka (in red) and no Eurêka (in blue) within the correct trials. Each dot represents a subject, color boxes represent the upper and lower quartiles, the black horizontal line within the boxes symbolizes the median, and the filled square is the mean value across subjects. **D**. Averaged RT of correct trials with Eurêka (in red) and without Eurêka (in blue). Same legend as in **C**. ns: non-significant, *: *p*<0.05.

On average, the participants reported a Eurêka in 55.6% (SD=20) of correct trials (and in 22.7% (SD=16.4) of incorrect trials), whereas they declared no Eurêka in 44.1% (SD=21.1) of correct trials (and in 49.1% (SD=20.5) of incorrect trials). Within correct trials, the percentages of Eurêka and no Eurêka did not statistically differ (W=149, *p*=0.26; **Figure 2C**). However, mean RT in trials correctly solved with Eurêka were significantly shorter than mean RT in trials solved without Eurêka (respectively, 6.8s (SD=2.1) and 11.4s (SD=3.8), W=15, *p*=1.30 10^−3^, **Figure 2D**).

We examined how *semantic remoteness* related to Eurêka reports by computing logistic regressions at the individual level (see method). The results show no significant effect of *SAS* (orthogonalized from RT) on Eurêka reports in any individual (**Figure S1A**). At the group level, the one-sample t-test of the individual regression coefficients was not significant (mean=0.004, SD=0.01, t(20)<1, *p*=0.70; **Figure S1B**).

### EEG

Time-frequency analyses were computed between 3 and 60Hz during the 2s period following the onset of the word triplet (initial time window) and the 2s period preceding the participant’s response (response time window; see method). Time-frequency maps were averaged along the frequency dimension according to four frequency bands (i.e., theta 3-7 Hz, alpha 8-12 Hz, beta 13-30 Hz, and gamma 31-60 Hz).

The average number of trials included in the EEG analyses (non-artifacted correct trials with RT>4s) for the initial and response time windows was, respectively, 32.1 (SD=11.5) and 32 trials (SD=11.7) for *semantic remoteness* condition, and 30.3 (SD=12) and 29.4 trials (SD=11) for the in *insight solving* condition. The time-frequency maps of EEG power across all trials are shown in **Figure S2** for each time window of interest, including topographical maps for each frequency band.

To explore the neurophysiological correlate of *semantic remoteness* and *insight solving*, we used a two-level statistical analysis approach. First, individual linear regressions assessed the relation between EEG power in each frequency band and behavior with EEG power as the dependent variable and i) semantic distance as the independent variable to explore the effect of *semantic remoteness*, ii) Eurêka self-report as the independent variable to explore *insight solving*. Then the resulting individual regression coefficients were tested at the group level (one-sample t-tests) with cluster-based corrections for multiple comparisons in spatial (65 electrodes) and time dimensions. These analyses were performed for each time window (see method).

Finally, we used source localization to explore the brain regions associated with the significant clusters observed at the sensor level. For this, we analyzed the cortical sources in the time windows and the frequency bands in which significant clusters were found.

#### Semantic remoteness in associative combination

We found three significant negative clusters (i.e., the lower the *SAS*, i.e., the more remote the solution, the higher the power in the considered frequency band) (**Figure 3**). No positive clusters were found.

**Figure 3.**
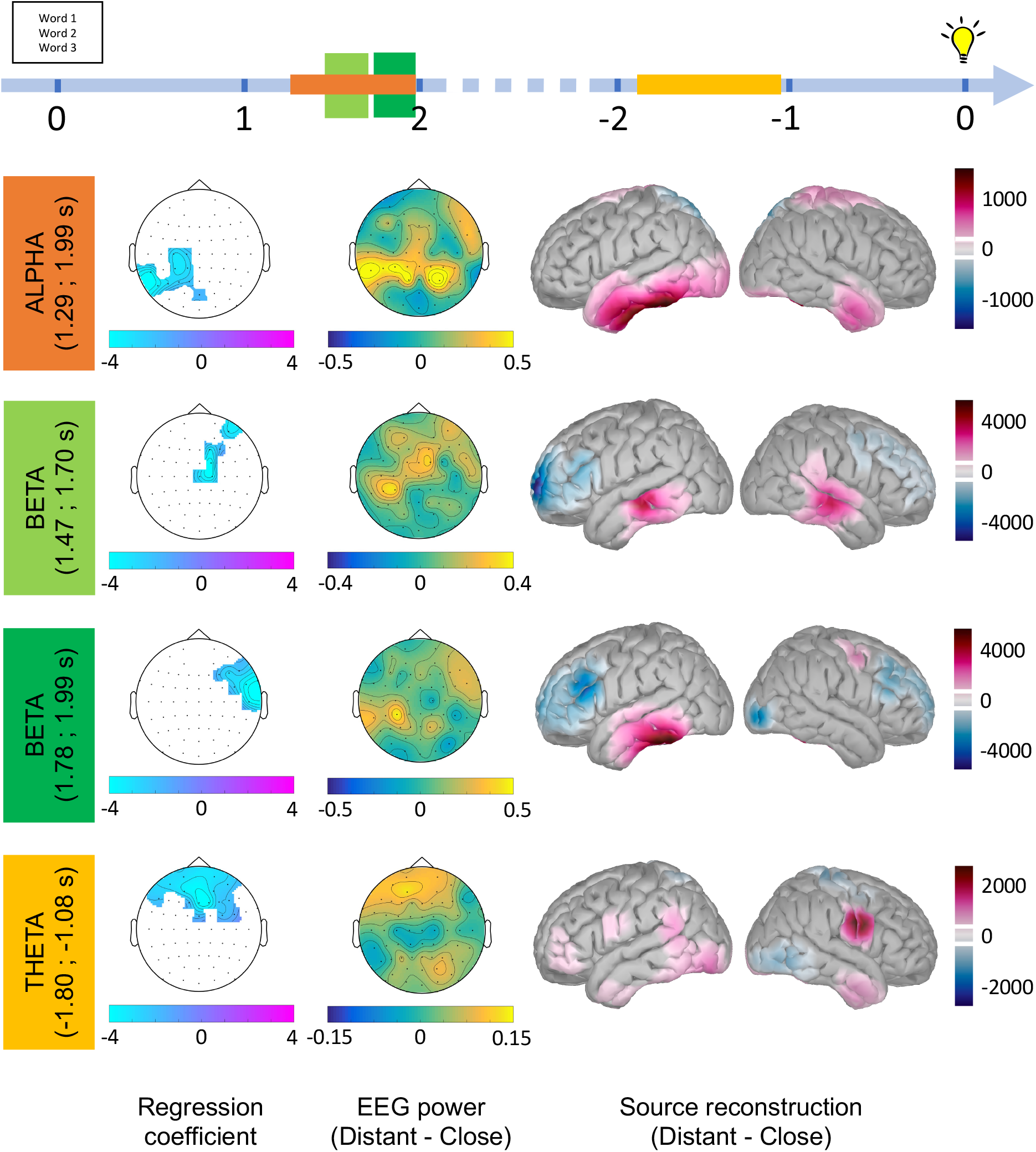
EEG effects related to the *remoteness* of semantic associations. **Top:** Time course (in second) of the task with the two time windows of interest (initial time window between 0 and 2s after the onset of the cue words, and response time window between −2 and 0 s before the response). Colored rectangles symbolize the time period where clusters significantly associated with the *remoteness* of semantic associations were observed in the alpha band (in orange), beta band (in light and dark green), and theta band (in yellow). For each cluster, the results are further detailed as follows. **First column:** Topographical maps of the clusters. The significant clusters (*p*_corr_<0.05) are represented for each frequency band. The color codes the regression coefficient values in the significant clusters (color bar from negative values in light blue to positive values in purple). **Second column:** Topographical maps of the EEG power in each band contrasted between distant (with low *SAS* values) minus close (with high *SAS*) trials, averaged across subjects and in the time-windows of the clusters (as indicated in the colored rectangles on the left). Color bars indicate EEG power (z-score of dB) from negative (in blue, distant<close) to positive (in yellow, distant>close) values. **Third column:** Source reconstruction of EEG activity in each frequency band during the significant cluster time periods. The cortical source maps were contrasted between conditions (distant - close), and we represent the difference in source activity, for each frequency, averaged in the time window of the cluster. The color bar indicates the power (in pA.m) from negative (in blue) to positive (in purple) values. The white lines in the color bars indicate the threshold used to visualize the source on the normalized cortical surface rendering.

Two significant clusters were observed during the initial time period. A first negative cluster was observed in the alpha band on left temporal and parietal electrodes, from 1.29 to 1.99s after the onset of the cue words (11 electrodes, sum(t)=-1523, *p*_corr_=5.99 10^−3^) (**Figure 3**, “alpha” in orange). We performed a source reconstruction of alpha band activity during the cluster time window and contrasted the cortical source maps between distant and close trials. The largest source differences in alpha band between 1.29 and 1.99s were located in the left inferior temporal gyrus and the left anterior part of the middle temporal gyrus. We also observed source differences in alpha band activity in the right hemisphere in the anterior part of the inferior and middle temporal gyrus and the right pre-and post-central gyrus (**Figure S3A**).

A second negative cluster was observed in the beta band during the initial time window. It was formed from two subsets of electrodes over time. Beta activity increased with *remoteness* first on central electrodes from 1.47 to 1.70s after the cue words onset (6 electrodes, sum(t)=-421, *p*_corr_=0.03) (**Figure 3**, “beta” in light green) and second on temporo-frontal electrodes from 1.78 to 1.99s (7 electrodes, sum(t)=-554, *p*_corr_=0.02) (**Figure 3**, “beta” in dark green). We performed source reconstruction in the beta band during these two time periods separately. Between 1.47 and 1.70 s, the distant versus close contrast revealed sources located in bilateral posterior middle temporal gyrus. In addition, there was reduced beta activity for distant than close trials in the left anterior part of the middle frontal gyrus (**Figure S3B**). Between 1.78 and 1.99s, the sources showing differentiated beta band activity for remote versus close trials were located in similar regions (**Figure S3C**): beta power was higher in distant than close trials in a potential source located in the left posterior inferior temporal gyrus and was lower in the left posterior and inferior gyrus encompassing the left inferior frontal sulcus.

The third negative cluster was observed in the theta band during the response time window (−1.80 to −1.06s before the response) on prefrontal electrodes (15 electrodes, sum(t)=-2574, *p*_corr_=2.00 10^−3^) (**Figure 3**, “theta” in yellow). As for the previous clusters, we reconstructed the sources of theta band activity in the cluster time period. Contrasting distant versus close trials revealed sources located in the right inferior part of pre- and post-central gyrus and in several regions in the left hemisphere, including the lateral part of the orbital gyrus and the anterior part of the inferior frontal gyrus, the inferior pre- and post-central gyrus, the posterior part of the superior temporal gyrus and posterior and anterior temporal areas (**Figure S3D**).

#### Insight problem-solving

We found three significant positive clusters, where trials solved with a Eurêka were associated with significantly higher activity amplitudes than those solved without a Eurêka. All clusters were observed in the response time window (**Figure 4**). No negative clusters were found.

**Figure 4.**
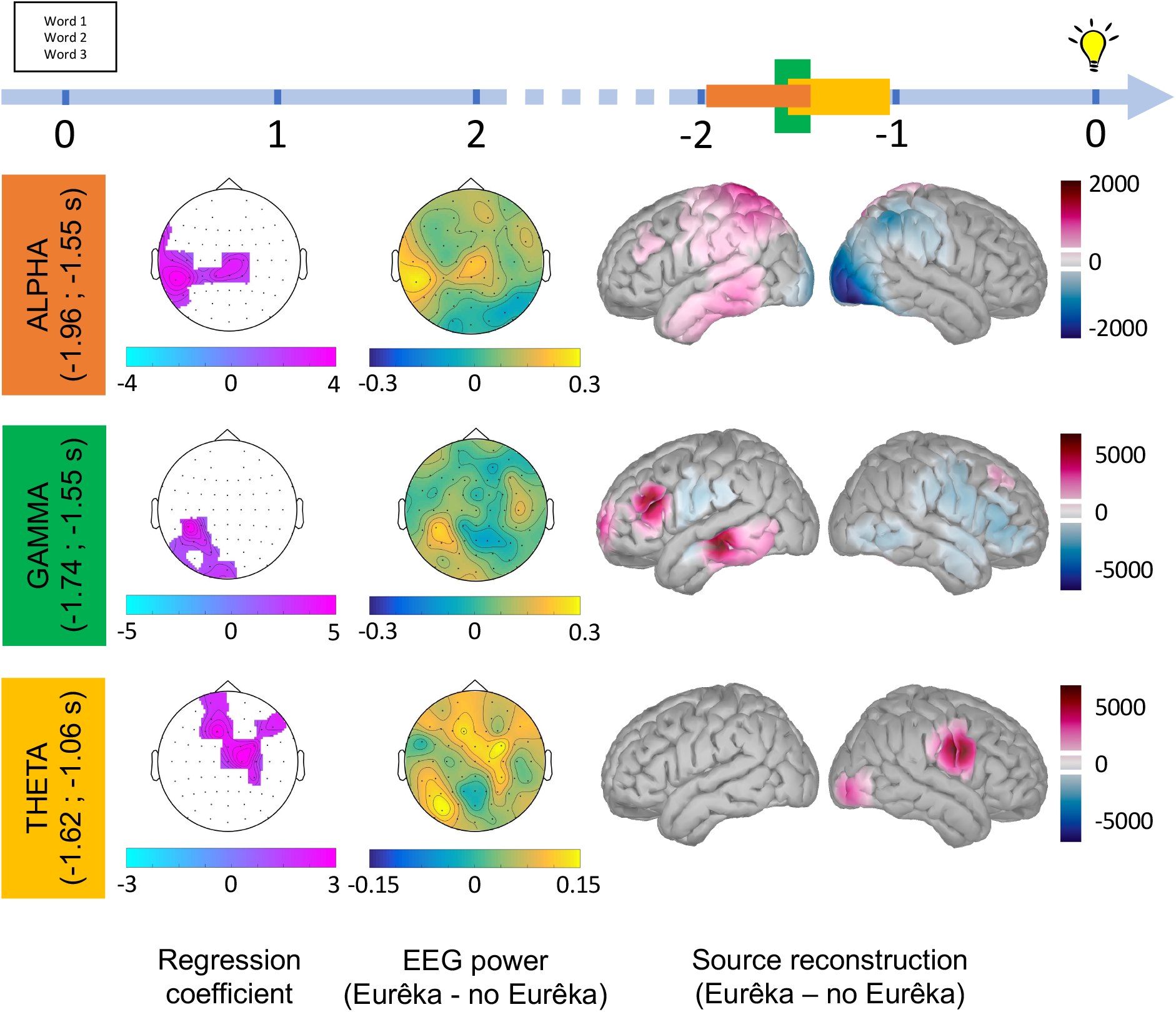
EEG effects related to *insight solving*. **Top:** Time course of EEG activity during the two time windows of interest (initial time window between 0 and 2s after the onset of the cue words, and response time window between −2 and 0s before the response). Colored rectangles symbolize the time period where clusters are significantly associated with Eurêka reports in the alpha band (in orange), the gamma band (in green), and the theta band (in yellow). For each cluster, the results are further detailed as follows. **First column:** Topographical maps of the cluster. The significant clusters (*p*_corr_<0.05) are represented for each frequency band. The color codes the regression coefficient values in the significant clusters (color bar from negative values in light blue to positive values in purple). **Second column:** Topographical maps of EEG power in each band contrasted between trials with Eurêka minus those without Eurêka, averaged across subjects and in the time windows of the clusters (as indicated in the colored rectangles on the left). Color bars indicate EEG power (z-score of dB) from negative (in blue, Eurêka<no Eurêka) to positive (in yellow, Eurêka>no Eurêka) values. **Third column:** Source reconstruction of EEG activity in each frequency band during the time periods of the significant cluster. The cortical source maps were contrasted between conditions (Eurêka – no Eurêka), and we represent the difference in source activity, for each frequency, averaged across the time window of the cluster. The color bar indicates the power (in pA.m) from negative (in blue) to positive (in purple) values. The white lines in the color bars indicate the threshold used to visualize the source on the normalized cortical surface rendering.

The first positive cluster was observed in the alpha band frequency in left central and temporal electrodes, between −1.96 and −1.55s before the response button press (13 electrodes, sum(t)=1174, *p*_corr_=0.01) (**Figure 4**, “alpha” in orange). The source reconstruction of EEG activity in the alpha band during the time window of this cluster showed increased alpha activity of sources mainly located in the left superior parietal lobule and posterior part of the inferior and middle temporal gyrus. Additionally, sources’ alpha activity was reduced in the right occipital polar cortex when participants reported a Eurêka (**Figure S4A**).

The second positive cluster overlapped temporally with the end of the first cluster during the response window (−1.74 to −1.55s before the response) and was found in the gamma band in left parieto-temporal electrodes (9 electrodes, sum(t)=372, *p*_corr_=7.99 10^−3^) (**Figure 4**, “gamma” in green). The source reconstruction of EEG activity in the gamma band during the time period of this cluster showed increased gamma activity for Eurêka relative to no Eurêka trials in the left anterior superior frontal gyrus, around the inferior frontal sulcus (encompassing posterior part of inferior and middle frontal gyrus) and left middle temporal gyrus (**Figure S4B**).

The last positive cluster was observed in the theta band on centro-frontal electrodes from −1.62 to −1.06s before the response (14 electrodes, sum(t)=1377, *p*_corr_=7.99 10^−3^) (**Figure 4**, “theta” in yellow). During the cluster time window, the source reconstruction of EEG activity in the theta band showed greater theta activity in the inferior part of the right pre- and post-central gyrus (**Figure S4C**).

## Discussion

We explored the neurophysiological correlates of two cognitive components of creative problem-solving. We used an adapted version of Mednick’s task (Mednick, 1962; Bendetowicz et al., 2017, 2018) to examine the time course of EEG power related to the *insight solving* and, for the first time, the effect of the *remoteness* of the solution to be found in the context of a semantic associative combination. In contrast with most of the previous EEG studies using a similar task that averaged signal across long time-windows or large set of electrodes, or were restricted to specific electrodes or frequency bands (Jung-Beeman et al., 2004; Razumnikova, 2007; Danko et al., 2009; Luft et al., 2018), we employed a data-driven time-frequency approach. We found distinct patterns of activity in several frequency bands associated with *remoteness* and *insight solving. Remoteness* was associated with a significant increase in alpha activity in a left temporo-central cluster and beta activity in a right fronto-temporal cluster during the initial phase of the task and a later increase in theta activity in a frontal cluster just before the response. EEG activity changes related to *insight* were observed uniquely in the period just preceding the response. They included an increase in alpha activity in a left temporo-central cluster, followed by a gamma activity increase in a left parietal cluster, and finally an increase in theta band activity in a fronto-central cluster. Overall, these EEG findings provide new insights into the dynamic mechanisms involved in creative problem-solving.

In the following sections, we discuss each result, first, at the sensor level (where robust two-level statistical analyses corrected for multiple comparisons allowed us to identify clusters with specific differences in several frequency bands of EEG activities), then at the source level (the brain areas that showed differences in activity in the frequency bands and time windows of sensor level clusters).

### *Remoteness* in associative combination

The *remoteness* of semantic associations was associated with an increase in activity in the alpha band, about 1.5 seconds after displaying the cue words. Alpha is the most reported EEG correlate in creativity studies using various tasks (Fink et al., 2009; Fink & Benedek, 2014; Fink & Neubauer, 2006; Jauk et al., 2012; Mölle et al., 1996; Shemyakina et al., 2007; Zhou et al., 2018; Mastria et al., 2021), including the RAT (Jung-Beeman et al., 2004; Sandkühler & Bhattacharya, 2008; Luft et al., 2018). Alpha activity increases with the creative requirements of the task (Fink & Benedek, 2014). It has been interpreted as an active inhibition (Klimesch et al., 2007; Klimesch, 2012) of external, non-relevant stimuli, allowing the increase of internal processing (Cooper et al., 2006; Cona et al., 2020) and internally-oriented attention (Fink & Benedek, 2014; Lustenberger et al., 2015). Klimesh (Klimesch, 2012) postulated that alpha-related inhibition is needed to explore and navigate in semantic memory, which is organized as a network. More precisely, access to remote knowledge may require that closely related, but not relevant memory information, is inhibited. Distant CAT trials likely required extended access to the knowledge stored in semantic memory as participants had to find a remote solution and inhibit close but irrelevant associations. The increase in alpha band activity during the initial time may reflect this process. The source reconstruction suggested that the effect of *remoteness* in the alpha band involved the left (and to a lesser extent to the right) inferior and middle temporal gyrus. Previous studies have identified different temporal regions as key brain areas for semantic processing (Hickok & Poeppel, 2004; Binder et al., 2009; Visser et al., 2012; Ralph et al., 2017) with distinct roles for regions along the rostro-caudal and supero-inferior axes (Ralph et al., 2017). The anterior temporal lobe appears as a transmodal hub in semantic processing in interaction with more posterior temporal areas. The left inferior temporal gyrus plays a role in semantic representation and word meaning (Whitney et al., 2011). The left posterior middle temporal gyrus is involved in a semantic control network (Noonan et al., 2013; Teige et al., 2019; Evans et al., 2020; Vatansever et al., 2021). Semantic control is likely involved in CAT, especially in the distant trials where participants had to retrieve remote associations and combine them. Neuroimaging studies using RAT-like tasks have reported the involvement regions of the semantic control network (Anderson et al., 2009; Gonen-Yaacovi et al., 2013; Jefferies & Wang, 2021). The involvement of semantic control in CAT is also consistent with previous research linking alpha activity with cognitive control (Sadaghiani & Kleinschmidt, 2016). Overall, the initial alpha activity that we found may reflect enhanced controlled access to the knowledge required by distant trials.

The *remoteness* of the associative combination was also associated with an early increase in beta power in the right centro-temporal and temporo-frontal electrodes. This beta activity temporally overlapped with the alpha cluster described above. Variation of beta activity during creative thinking or problem-solving is not classically reported. A few studies reported an increase in beta activity in frontal and temporal electrodes associated with the RAT (Razumnikova, 2007) or during other creativity tasks (Rosen & Reiner, 2016; Zioga et al., 2020), but its functional role in the context of creativity is not understood. Beta activity is usually associated with motor preparation, but in different regions and time windows than in our study (da Silva, 2009; Weiss & Mueller, 2012). Hence, the observed higher beta activity for more remote combinations may reflect non-motor cognitive processes. Enhancement of beta-band activity has been related to various aspects of language processing (Weiss & Mueller, 2012), such as the maintenance of a mental state during a cognitive task requiring language (Engel & Fries, 2010) or of visual object representation in short-term memory (Tallon-Baudry et al., 1999). The role of beta activity increase in distant CAT-solving is not obvious. One can speculate that when the cue words to be combined are not quickly converging to a solution, the current mental activity (i.e., active exploration of semantic memory related to the alpha activity) should be maintained, increasing beta activity in the distant condition.

Finally, *remoteness* was associated with higher theta activity one second before the subject’s response, involving fronto-temporal regions. Theta activity in creative problem-solving has been scarcely reported (Razumnikova, 2007; Sandkühler & Bhattacharya, 2008). The role of theta activity in cognition is debated. Prefrontal theta activity has been associated with several aspects of executive control functions. Cavanagh and colleagues (Cavanagh et al., 2012; Cavanagh & Frank, 2014) proposed that theta rhythm generated by the median PFC region is involved in monitoring novelty, conflict, and surprise. Theta activity increases when information is accumulated (Cavanagh et al., 2012; Cavanagh & Frank, 2014). When controlled processes are engaged during goal-directed behavior, theta band coherence between frontal and other relevant brain regions increases (Zavala et al., 2018). Several studies have also associated theta activity with other controlled processes and functions such as inhibition (Adelhöfer & Beste, 2020), planning (Domic-Siede et al., 2020), prioritizing relevant information in working memory (Riddle et al., 2020), or analytical reasoning (Williams et al., 2019). Importantly, theta activity has been related to memory retrieval and encoding (Düzel et al., 2010) and may reflect integration processes that allow us to build new connections between elements of knowledge in semantic memory (Backus et al., 2016; Nicolás et al., 2021). Hence, theta band activity associated with distant CAT may reflect controlled retrieval and integration in semantic memory. Consistent with this interpretation, the central contribution of executive and memory processes in creativity is now well established (Cassotti et al., 2016; Beaty et al., 2016; Volle, 2017; Benedek & Jauk, 2018; Benedek & Fink, 2019). Recent studies have demonstrated the important role of the executive control network for creative thinking (Beaty et al., 2016, 2017; Bendetowicz et al., 2018). The executive control network supports several control processes involved in creative thinking such as working memory, inhibition, attentional control, planning, flexibility, and control and selection in memory retrieval. The source reconstruction of our theta-related cluster revealed a set of left regions largely coherent with the executive control network, such as the rostro-lateral PFC, parieto-temporal junction, and temporal regions. The rostro-lateral part of the PFC is a node of the executive control network that has been shown critical for solving CAT in frontal patients, especially in distant trials (Bendetowicz et al., 2018). Additionally, the grey matter volume in this region was also correlated with performance at this task (Bendetowicz et al., 2017). The particular role of the left rostro-lateral PFC in the CAT may be to combine the retrieved associates or integrate the result of the search from each cue word, i.e., in the relational integration of distant items (Aichelburg et al., 2016; Green et al., 2016; Urbanski et al., 2016). Thus, observing theta power increase during the response time window is consistent with previous studies using different methods showing the involvement of the left rostro-lateral PFC and executive control network in creativity. The source reconstruction also located theta activity in the right inferior pre- and post-central gyrus, a result that was shared between *remoteness* and *insight* analyses and is discussed below.

Overall, our results combined with the existing literature suggest that remote associative semantic combination relied on several controlled processes in distinct periods of CAT-solving (**Figure 5**). Hypothetically, in the initial phase of the task, semantic control (supported by alpha activity in the posterior middle temporal gyrus) may enable the exploration of semantic memory in search of remote associates (in relation to alpha activity in infero-temporal regions, including the temporal pole). The semantic search or search space might be reflected in the overlapping beta activity, that is associated with the maintenance of a current mental state or representation. Finally, just before the response, the increased prefrontal theta activity may reflect the involvement of other executive controlled processes, allowing to integrate and combine the search results from each cue word, evaluate the generated candidate solution, and finally select the most appropriate response.

**Figure 5.**
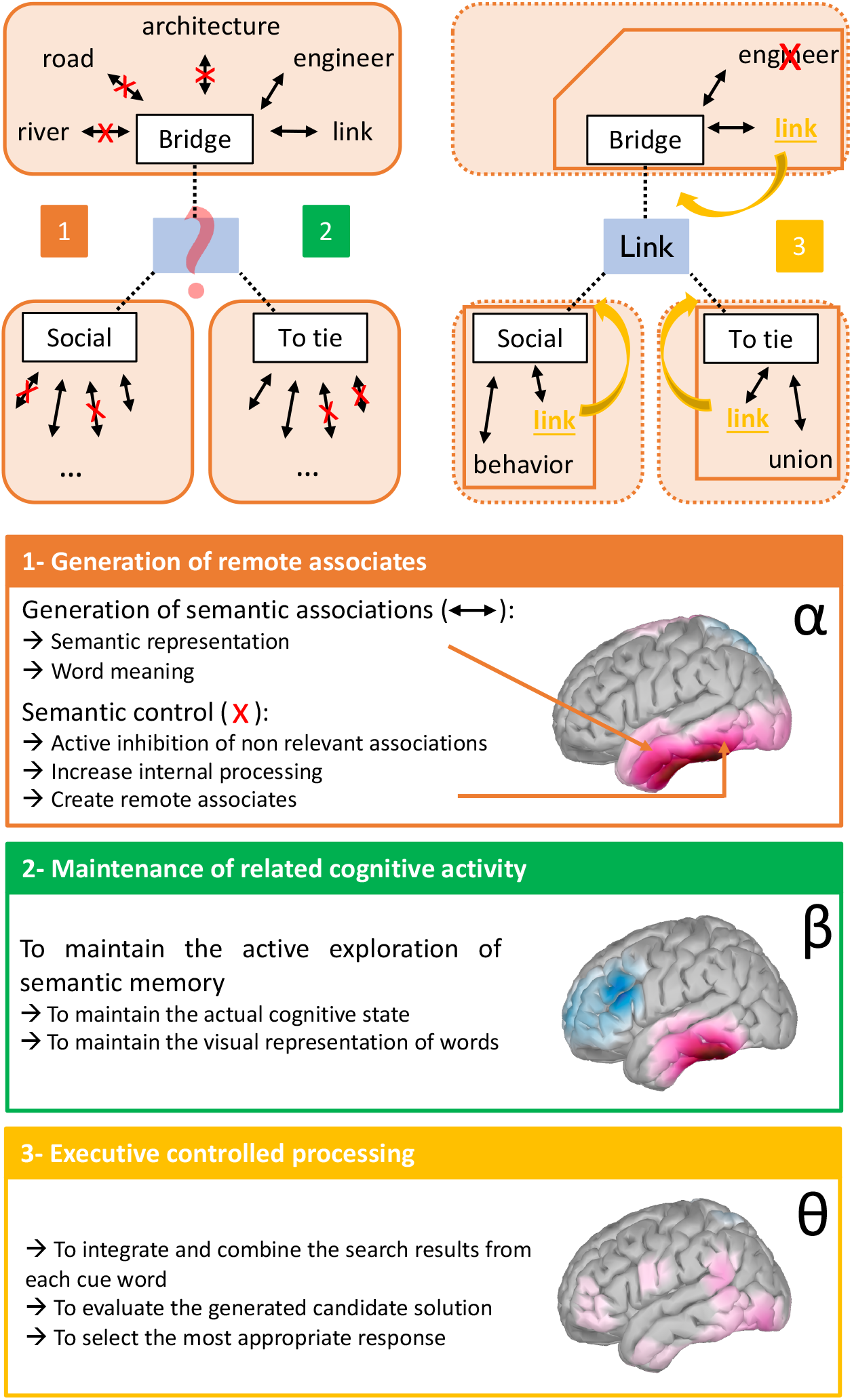
Hypothetical model of remote associative combination. Combined with the existing literature, our results suggest that solving a distant CAT (relative to a close one) requires generating remote semantic associates to each of the three cue words. This is supported by alpha activity in temporal areas (in orange). Overlapping beta activity might facilitate this process by maintaining related cognitive activity (in green). Then, executive controlled processing is needed to integrate, combine, evaluate and select the appropriate response. This final step is supported by theta activity found in brain regions involved in the executive control network.

#### Insight solving

The second aspect of the CAT that we analyzed, the effect of *insight solving*, was associated with distinct EEG correlates than remote associative combinations. Eurêka-related EEG differences were observed only during the response time window, suggesting that the early stages of problem-solving were similar for trials with or without Eurêka. It might be explained by the fact that the two solving modes (with and without Eurêka) are not exclusive and may co-occur within a trial. It is possible that people initially used analytical thinking until they reach an impasse and finally solved the problem with insight. Cognitive theories link insight with the need to experience a mental impasse and to restructure the problem representation before solving it with insight (Ohlsson, 1992; Sandkühler & Bhattacharya, 2008). It may thus be not surprising that insight and non-insight trials only differ in the period just preceding the response. Nevertheless, we examined only the first and last two seconds of problem-solving. We cannot exclude that differences between trials with Eurêka and without Eurêka occurred in between these time windows.

Just before the response, we observed successive modulation of alpha- and gamma-band activities for trials with Eurêka (compared to those without Eurêka), which is consistent with previous EEG studies (Jung-Beeman et al., 2004; Sandkühler & Bhattacharya, 2008; Sheth et al., 2009; Oh et al., 2020). The source reconstruction suggested that alpha activity related to Eurêka involved the left posterior inferior and middle temporal gyrus and the parietal region. Although the alpha activity associated with the *remoteness* and *insight solving* effects showed some similarities, they occurred at different time periods (initial vs. response time windows), suggesting that they reflected distinct mechanisms. Given the role of alpha in inhibition processes and the involvement of the left inferior and middle temporal gyrus in semantic processing discussed above, the Eurêka-related alpha increase may reflect the inhibition of non-relevant information to overcome the mental impasse and restructure the problem (Sandkühler & Bhattacharya, 2008). The source reconstruction also suggested that the superior parietal lobule, a region often showing alpha activity in relation to creativity (Fink & Benedek, 2014), played a role in *insight solving*. Given the classical role attributed to alpha activity in parietal areas in creativity research, this cluster might alternatively or additionally reflect in increased state of internally oriented attention during trials with *insight solving* (Fink & Benedek, 2014).

Succeeding to alpha, we observed a gamma activity increase in left parietal electrodes, which involved the left anterior superior frontal gyrus, left posterior inferior, and middle frontal gyrus and left middle temporal gyrus. An increase in gamma activity is often reported by studies exploring insight problem-solving (Jung-Beeman et al., 2004; Sandkühler & Bhattacharya, 2008; Sheth et al., 2009; Rosen & Reiner, 2016; Oh et al., 2020) and was related to the suddenness of the solution (Sandkühler & Bhattacharya, 2008). Gamma burst has been associated with the sudden awareness of a mental representation from memory (Tallon-Baudry et al., 1999; Engel et al., 2001; Engel & Singer, 2001). Hence, the gamma activity observed during the CAT-solving with *insight* may reflect the awareness of a solution that popped up suddenly in mind, yielding the subjective Eurêka experience.

The similar alpha followed by gamma synchronization associated with insight reported by previous studies involved distinct electrodes that we observed, especially in the right hemisphere (Jung-Beeman et al., 2004; Sandkühler & Bhattacharya, 2008; Sheth et al., 2009). The reasons for this left-right difference with our results are unclear. They might relate to the use of different paradigms. Previous studies mostly used the compound remote associate task, requiring finding a word that forms a compound word with each cue. Instead, we use a version where the solution is associatively related to the cue words. Thus, our task may rely more on semantic processing than the compound remote associate task, thus recruiting more left-brain areas (Hickok & Poeppel, 2004; Gonzalez Alam et al., 2019). Another methodological difference is that previous EEG studies on insight focused on specific scalp regions or frequency bands based on a priori hypotheses. In contrast, we used a data-driven approach considering all the electrodes and frequencies in our analyses while controlling for multiple comparisons. It potentially revealed new brain correlates of insight problem-solving. In addition, as in other EEG studies based on the RAT (Sandkühler & Bhattacharya, 2008; Oh et al., 2020), we observed alpha and gamma effects earlier than in Jung Beeman and al study (Jung-Beeman et al., 2004). This difference may relate to the instructions given to our participants of pressing the space bar when they thought the solution they had in mind was correct. It may have encouraged the participants to evaluate their solution more carefully and added a delay between the insight moment and button press.

Finally, as for remote trials, insight trials were associated with higher theta activity in frontal electrodes. This theta activity may reflect conflict monitoring because when the solution arises suddenly in consciousness, a conflict (or surprise) with the ongoing mental representations or ideas can arise, signaling a need for monitoring and selection. The source reconstruction located a potential source in the right inferior part of the pre- and post-central gyri. Although sensorimotor regions in creativity has already been described (Matheson & Kenett, 2020), its role remains challenging to interpret. Interestingly, this region was also a candidate source for the theta activity associated with *remoteness* during the same time window. *Remoteness* in associative combination and *insight solving* are often confused in previous studies (Dietrich & Kanso, 2010), leading to an unclear link between them. According to some theory, they are both resulting in overcoming a mental impasse suggesting that they might share similar thread in the time course of EEG power. In our study, we explored both components with the same task. Overall, our results did not suggest a link between *insight solving* and *remoteness* (no significant interaction at the behavioral level, distinct brain cluster at the sensor level). The shared sources in the theta band frequency cannot be explained by an imbalance in the distribution of trials between the two conditions (for instance, more Eurêka reports in distant trials) as the average number of trials included in the EEG analyses did not significantly differ between conditions (see *Supplementary Data*). Even if our source reconstruction is not specific to the cluster found at the sensor level (but rather to a frequency band during a specific time window), we cannot exclude that *remoteness* and *insight* effects shared some similar brain event occurring just before the response. Further study will be needed to clarify this question.

To summarize (**Figure 6**), we show that *insight solving* is associated with successively increased alpha, gamma, and theta power during the last seconds of a CAT-solving. Alpha activity could help to overcome strong but obvious associations of ideas. The solution could hence suddenly emerge in the individual’s mental representation, and lead to a gamma activity. Then, a conflict might occur between the Eureka-mediated solution and the previously ongoing mental thinking. This conflict needs to be monitored and controlled, which may be reflected by the increase in theta activity.

**Figure 6.**
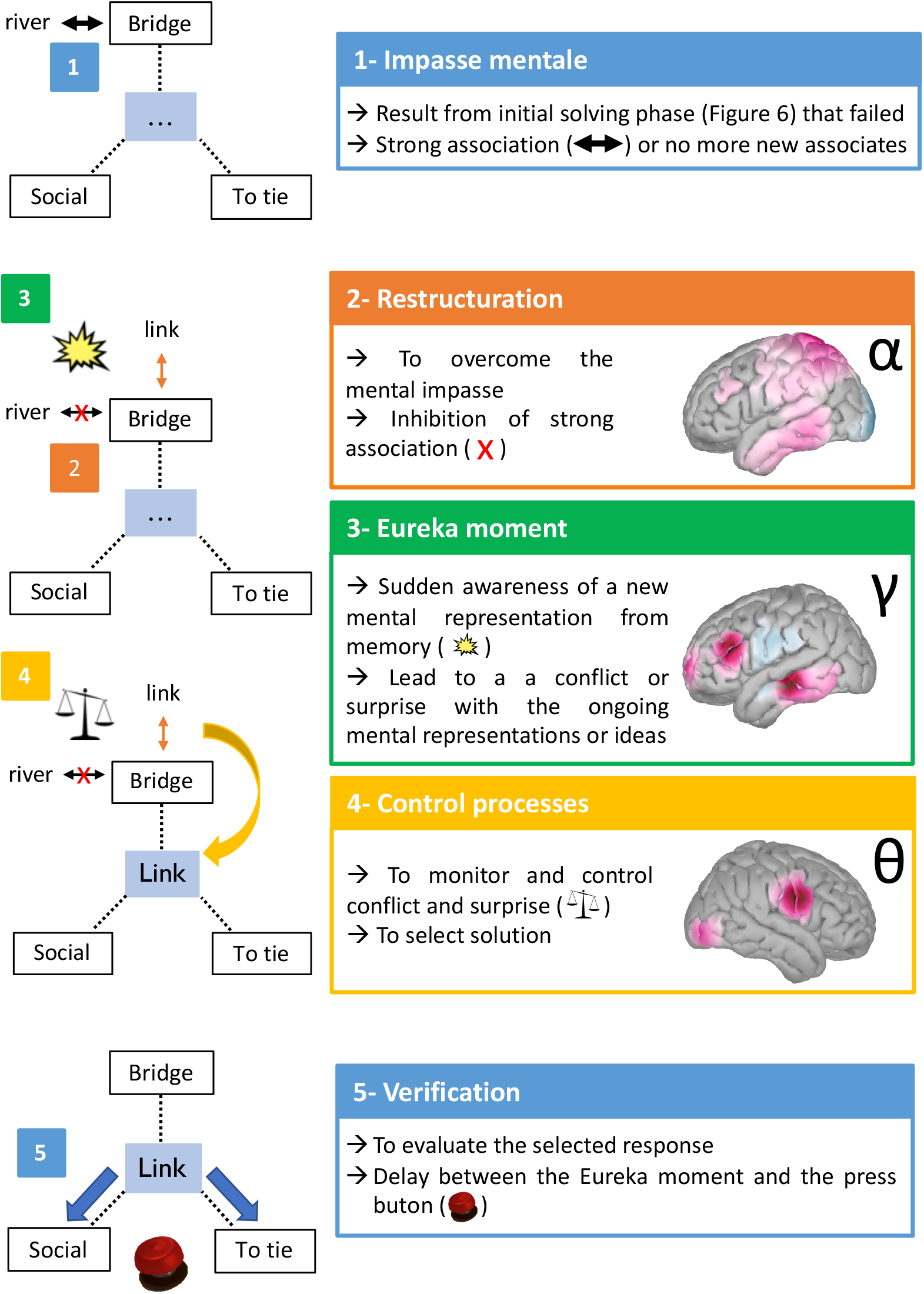
Hypothetical model of *insight solving* related results. After reaching a mental impasse (in blue), restructuration is needed to break out it. Inhibition of stong associations can be supported by alpha activity in temporal areas (in orange). When a new association suddenly arises in consciousness, a realization of the solution occurs (supported by gamma activity, in green). It ensures a conflict or surprise that needs to be monitored to select the appropriate association (supported by theta activity, in yellow), verified before answering (in blue).

## Limitations

This study is not without limitations. First, the CAT is a difficult task with a low correct response rate, often around 60%. Added to the constraints related to EEG artefact cleaning, we included in our analyses much fewer trials than expected. We chose not to analyze incorrect trials since the cognitive involvement of the participants in incorrect trials is uncontrolled. Second, the usually long response time in such a task led us to analyze fixed time windows at the beginning and end of each trial without considering the time in between. We thus do not provide the whole picture of the processes happening during our task. Finally, it may be noted that the individual MRIs of the participants were not available. Thus, source reconstruction results must be interpreted cautiously and entail more uncertainty than the effects we characterized at the sensor level. However, the involved regions are broadly consistent with the neuroimaging literature, and our results offer new perspectives on potential networks involved in creative problem-solving. Finally, we used the most used and validated approach with self-reports of Eureka experience to define *insight solving* (Laukkonen & Tangen, 2018). However, the best method to capture the insight phenomenon that would best reflect specific solving mechanisms is an open question.

## Conclusion

This study explored the EEG correlates of two aspects of RAT problem-solving, *remoteness* in associative combination and *insight solv*ing. We showed distinct patterns of brain activity in the time-frequency domain for these two aspects. First, *semantic remoteness* was associated with an early alpha and beta activity in latero-temporal regions and a theta activity in frontal areas just before the response. These results suggest that early controlled processes may guide and constrain the search of remote associates, whereas later controlled processes may integrate or combine the retrieved information. Second, *insight solving* was associated with alpha then gamma activity in infero-temporal regions and theta activity in frontal areas, which occurred just before the response. These findings indicate that *insight* is supported by specific brain dynamics distributed in space and time that may relate to a sudden restructuration of the problem or its solution. Furthermore, late theta activity might also suggest that solving a problem with *insight* also includes the involvement of control processes, possibly in the facilitation or monitoring of the Eurêka-mediated solution. Further work is needed to overcome approximates of source reconstructions. Combining neuroimaging approaches or recording intracranial EEG signal can be promising methods for future research to better understand brain correlates of creative problem-solving.

## Method

### Participants

Twenty-three right-handed native French speakers aged from 21 to 25 years old (mean age=23.04; standard deviation, SD=1.15; 13 women) were included in the study. All participants were healthy adults with MMSE ≥ 28 (Folstein et al., 1975), no history of neurological and/or psychiatric illness, no psychoactive substance abuse, nor consumption less than 24 hours before the experiment. Two participants were excluded because of technical problems during the experiment. The analyzed sample thus consisted of 21 healthy adults (mean age=22.95, SD=1.15 years old, 12 women). A national ethical committee approved the study. All the participants gave their written informed consent and received financial compensation.

### Experimental task

EEG was recorded during the performance of the CAT (Bendetowicz et al., 2017, 2018), which is an adapted version of the RAT (Mednick, 1962). In such a task, subjects are asked to provide a word that connects three unrelated cue words. Our adapted version varied the semantic strength association (*SAS*) between the cue words and the expected solution, based on French associative norms (Debrenne, 2011; Bendetowicz et al., 2017, 2018). We considered the average *SAS* between the expected solution and each of the three cue words within each trial. Hence, every trial was characterized by a *SAS* value: the lower the *SAS* value, the more remote the solution was from the cue words (an example trial with a low *SAS* value - hence distant solution - is Bridge-Social-To tie, where the solution is Link; an example trial with a high *SAS* value – hence close solution – is Street-Countryside-Centre, where the solution is Town). Previous studies using the CAT (Bendetowicz et al., 2017, 2018) have shown that the performance in this task, especially for distant trials, correlated with other creative assessments suggesting its external validity. Following the same principles as in the original CAT, we built 28 additional trials for the current study in order to anticipate the loss of analyzable trials due to EEG experimental constraints and artifacts. In total, each participant performed 100 trials (median *SAS* value 6.5, range from 0.3 to 38.8).

During the experiment, the participants were seated comfortably in front of a computer screen. Before starting the task, the examiner explained the general design and instructions with written support. Explanations on Eurêka were particularly detailed. It was described as “the subjective experience you can have when you solve a problem, and the solution comes to mind suddenly, it is not the result of cognitive efforts, and you are not able to report the mental steps leading to this solution”. It was opposed to analytic solving in which “you have a strategy and the feeling of gradually getting closer to the solution”. We clarified that these two solving methods were not incompatible or exclusive and instructed the participants to consider only a few seconds before their response. To ensure the participants understood the instructions correctly, they completed ten practice trials, and the instructions were repeated when needed. After instructions and training, the participants performed 100 trials in random order while EEG was recorded. Breaks were proposed to the participants every 25 trials to limit fatigue.

The CAT was computerized and programmed using the Psychtoolbox (version 3.0.11) running in MATLAB (MATLAB version 9.0 (R2016a), Natick, Massachusetts: The MathWorks Inc.)) (**Figure 1**). For each trial, the three cue words were displayed on the center of the screen, one above the other to limit eye movements as much as possible. The participants were asked to give a unique word related to all three cue words and had up to 30s to respond. They were aware that the response could be a noun, a verb, or an adjective but not a proper noun or a compound word. As soon as they thought they had found the correct answer, they pressed the space button of the keyboard. This made the three words disappear, and the participants had then a fixed time of 2.5s to tell their response verbally. The screen remained blank during this period. The examiner wrote down the participant’s response. In addition, as classically performed in previous studies using similar tasks, we collected the self-report “Eurêka” experience on a trial-by-trial basis (Kounios & Beeman, 2014). Thus, after the 2.5s response period, the question “Eurêka?” was displayed on the screen. The participants had to indicate whether the solution they gave came to their mind with a Eurêka by pressing the keyboard letters “V” (Eurêka) or “N” (no Eurêka) within a time limit of 5s. A central fixation cross was displayed during the intertrial interval followed by a jittered duration (mean=1.5s, range between 1.2s and 1.8s).

### Behavioral measures and analyses

Accuracy (correct or incorrect) was determined based on the French associative norm (Debrenne, 2011; Bendetowicz et al., 2017). Responses were also considered valid if they were lexically similar or synonyms to the one defined by the French associative norm. Finally, few additional answers were accepted if they provided semantic similarities with the cue words but were not in the French associative norm. In this case, only responses selected by a panel of five external judges were considered correct. We defined response time (RT) as the time between the onset of the display of the cue words and the space bar press.

Each trial is characterized by a *SAS* value (a continuous variable determined by the material and fixed between subjects) and can be categorized according to how the subject solved it (with or without Eurêka; binary variable that depends on each subject). To estimate the effect of *remoteness* (*SAS*) on performance, we computed the percentage of individuals with correct responses (i.e., number of individuals who gave a correct response divided by the total number of participants) and the mean RT for correct responses on a trial-by-trial basis. We explored the relation of accuracy and RT with the corresponding *SAS* value using Spearman correlations.

We also explored how many trials were solved (or not) with a Eurêka and without a Eurêka. To examine whether trials solved with a Eurêka differed from those without a Eurêka, we compared the averaged percentage of Eurêka and no Eurêka, and the averaged RT of trials with and without Eurêka across individuals. We focused on correct trials as incorrect ones were excluded from the EEG analysis. Statistical comparisons were performed using non-parametric paired Wilcoxon tests.

Finally, we explored the link between the effect of *SAS* and Eurêka using a two-level modeling approach. First, we ran a Global Linear Model (GLM; using the glmfit function in MATLAB) at the individual level using only correct trials. Taking advantage of the *SAS* value of each trial, we used logistic regression to explore whether the *SAS* predicted a Eurêka. As we expected the *SAS* to be correlated with RT, we removed the variance explained by RT from the *SAS* variable. We then computed a logistic regression exploring the relationship between the corrected *SAS* and Eurêka. Then, for the second-level analysis, we computed a one-sample two-tailed t-test (against zero) on the subject’s regression coefficients resulting from the GLM. This allowed us to analyze the relation between Eurêka reports and *SAS* at the group level.

### EEG

#### EEG recording

EEG data were recorded using BRAINAMP DC system (Brain Products GmbH, Münich, Germany) with 64-active electrodes mounted in an elastic cap (actiCAP) according to the extended International 10–20 system and including a row of low fronto-temporo-occipital electrodes (PO9/10, TP9/10, FP9/10). Two additional electrodes were used as reference (FCz electrode) and ground (AFz electrode). Disposable electrodes placed above and below the right or left eye and lateral to the outer canthus of both eyes recorded vertical and horizontal EOG, respectively. Electrode impedances were at or below ten kOhm. The EEG data were recorded at 1 kHz with an online 0.016-250 Hz bandpass filter.

#### EEG preprocessing

All EEG preprocessing and analyses were performed using the FieldTrip toolbox (Oostenveld et al., 2011), completed by homemade scripts, and brainstorm (version 09-Sep-2020) (Tadel et al., 2011) running under MATLAB (MATLAB version 9.0 (R2016a), Natick, Massachusetts: The MathWorks Inc.).

EEG signal was downsampled offline to 128 Hz, and filtered with zero phase, third order high pass, and low pass Butterworth filters (set at 0.5 and 63 Hz, respectively). Independent component analysis (ICA) was used to detect and remove artefacts caused by eye blinks. On average, two independent components (IC) were removed after the visual inspection of the time series and topographies of the IC. Then, the EEG signal was visually inspected to exclude artifacts related to muscles or movements. Next, noisy channels were interpolated using the averaged signal of adjacent channels. A mean of 7 electrodes (SD=2.1) was interpolated across participants. Trials containing more than 10% of bad channels were removed (11 trials per individual on average, SD=6.5). Finally, the signal was re-referenced to the average of all electrodes (recovering the FCz channel).

We segmented the EEG signal for each trial in two time windows of interest. First, the “initial time window” corresponded to the 2s period following the onset of the cue word display on the screen. Second, the “response time window” corresponded to the 2s period preceding the space bar press (i.e., the subject’s response). We considered only correct trials for EEG data analysis. We excluded the trials with an RT shorter than 4s to avoid overlapping our two time windows (14 trials excluded on average per individual, SD=9.5).

Averaged numbers of analyzed trials across individuals are presented in **Table S1**. In addition, supplementary analyses are provided to ensure there was no unbalance between the number of trials analyzed across conditions (i.e., *semantic remoteness* and *insight solving*; see supplementary material).

#### Time-frequency computation

Time-frequency maps were computed for each electrode, trial, and time window (initial and response) in a frequency range between 3 and 60 Hz. We used a multitaper time-frequency transform (Slepian tapers, lower frequency range: 3-32 Hz, six cycles, and three tapers per window; higher frequency range: 32-60 Hz, fixed time-windows of 240ms, 4-31 tapers per window). This approach allows better control of time and frequency smoothing. It uses a constant number of cycles across frequencies up to 32 Hz (hence a time window with a duration that decreases when frequency increases) and a fixed time window with an increasing number of tapers above 32 Hz in order to obtain more precise power estimates by adaptively increasing smoothing at high frequencies. Hence, the resulting EEG power represents the signal amplitude in a given frequency after its spectral decomposition. Time courses were aligned to the onset of the cue word display for the initial time window (corresponding to time 0 for the initial time window epochs) and the space bar press for the response time window (corresponding to time 0 for these latter epochs). We performed a z-score baseline correction of time-frequency maps using the time-frequency maps computed from the EEG signal recorded −1.2 s to −0.1 before the onset of the display of the cue words on each trial. Finally, time-frequency maps were averaged along the frequency dimension according to the four frequency bands: theta 3-7 Hz, alpha 8-12 Hz, beta 13-30 Hz, and gamma 31-60 Hz.

#### Task-based analysis

As for the behavioral analysis, we used a two-level statistical analysis approach at the sensor level. First, we used individual linear regressions to explore the relation between EEG power and behavior. To explore EEG correlates of *semantic remoteness*, we used EEG power as the dependent variable and *SAS* as the independent variable. To explore *insight solving*, EEG power was the dependent variable, and the Eurêka report was the independent variable. These two analyses were performed independently at the individual level for each point in time, in each frequency band (theta, alpha, beta, gamma), and for each time window (initial and response time window). This first level of analysis allowed us to obtain regression coefficients at the individual level. Then, at the second (group) level, the resulting individual regression coefficients were analyzed at the between-subject level with a one-sample two-tailed t-test against zero. According to the following procedure, we corrected our results for multiple comparisons using a cluster-based correction for the time and space (electrode) dimensions. For each frequency band and time window of interest, the results from the one-sample t-tests performed at each time point and on each electrode were clustered based on spatio-temporal and statistical criteria. The cluster spatial extent was defined as at least one neighboring electrode in either time or space based on the template “easycapM1” provided by the Fieldtrip toolbox and matched our electrode cap. The clusters formation considered only the (time, electrode) points where the one-sample t-tests were significant with a *p-value* lower than 0.0125. We selected this statistical threshold because we computed a cluster-based analysis for each of the four frequency bands of interest (0.05/4=0.0125). Then, we computed the sum of the t-test statistics within each obtained cluster (sum(t)). In order to obtain the distribution of this cluster statistics under the null hypothesis while correcting for multiple comparisons, we repeated this analysis on 1000 Monte Carlo randomizations, retaining only the maximum value of the sum of t-test across clusters on each randomization. The clusters obtained from the original data were finally considered significant if their *p*-value (*p*_corr_) was lower than 0.05 across the 1000 randomizations.

#### Source reconstruction

We explored the brain regions related to the significant clusters observed at the sensor level using source localization. For this, we analyzed the cortical sources in the time windows and the frequency bands in which significant clusters were found. We used the Brainstorm software that is freely available for download online under the General Public License (http://neuroimage.usc.edu; (Tadel et al., 2011)).

For each individual, first, a head model was computed using the symmetric boundary element method (BEM) method from OpenMEEG open-source software (Gramfort et al., 2010), based on the template MRI normalized in the Montreal Neurological Institute (MNI) system, available in Brainstorm software (MNI/Colin27), and coregistered with the 65 electrodes considering standard 10-10 electrode coordinates. Next, the noise covariance matrix was computed on the time window of interest of all trials with a baseline corresponding to the time period preceding the onset of the word triplet (−1.2s to −0.1s). Sources were then computed at the trial level using preprocessed EEG signal (that is, 128-Hz, ICA-corrected, average-referenced EEG signal). Next, we applied a weighted minimum norm imaging (wMNE) method with current density map measures computed for 15000 trihedral dipoles – total of 45000 elementary dipoles, equivalent to sources unconstrained in their orientation – distributed over the cortical mantle of the brain model obtained from the standard MNI/Colin 27 brain template. Then, we computed the power within the considered frequency band using a Hilbert transformation at the source level for each cluster identified at the sensor level (i.e., in each time window and frequency band of interest). Since we used unconstrained orientations for the sources, we computed the time-frequency decompositions for all 45000 elementary dipoles and summed the power for the three orientations at each source location (or vertex) as recommended. Finally, power was averaged within the time window of the cluster and then averaged across trials separately for each studied experimental condition. This procedure was repeated for each participant, and the obtained cortical current power maps were averaged across participants in each condition. Then, we contrasted the maps between conditions (Distant minus Close conditions or Eurêka minus no Eurêka conditions) according to the considered cluster. We did not run further statistical analysis at the source level to avoid double-dipping. The cortical current power maps were thresholded to visualize only sources with activity higher and lower than 10% of the absolute maximal source.

## Acknowledgements

We thank all the participants to the study. EV and TB are funded by the ‘Agence Nationale de la Recherche’ [grant numbers ANR-19-CE37-001-01], the ‘Fondation pour la recherche medicale’ [grant number DEQ20150331725]. The research also received funding from the program ‘Investissements d’avenir’ ANR-10-IAIHU-06. MOT is funded by Becas-Chile of ANID (CONICYT). TB is funded by ‘Société Française de neurologie’ and AP-HP. NG and LH are funded by the program “Investissements d’avenir” (Agence Nationale de la Recherche, grant numbers ANR-10-IAIHU-06 and ANR-11-INBS-006) for infrastructure funding. BG received grant number from the ‘Fondation pour la recherche médicale’ [FDM20150632801].

## Authors’contribution

Conceptualization: EV and TB; Methodology: TB, MOT, BG, LH, NG, KL and RL; Data acquirement: TB, MOT and BG; Analyses: TB, ALP, NG and KL; Supervision: EV and NG; Writing: TB, EV and NG.

## Data availability

The material and the data sets generated during and/or analyzed during the current study are available from the corresponding author (TB) on reasonable request.

