## Supplementary data for "Time course of EEG power during creative problem-solving with insight or remote thinking"

### Supplementary file

#### Supplementary materiel

To test if there was a difference between the number of trials analysed between the conditions related to the *remoteness* (high vs. low *SAS* values) and *insight solving* (with vs. without Eurêka), we performed a 2x2 ANOVA. For this purpose, we categorized trials according to their *SAS* value into distant trials (with *SAS* value lower than the overall median *SAS* value) and close trials (with a *SAS* value greater than the median *SAS* value). **Table S1** indicates the average number of analyzed trials in each condition.

In line with the previous results, there was a significant effect of *remoteness* on the distribution of analyzed trials in both time windows (initial time window: $F(1,80)=5.83$, $p=0.02$; response time window: $F(1,80)=4.65$, $p=0.03$), meaning that the number of analyzed trials was greater in the close than the distant condition ($p<0.05$). There was no significant difference in the number of analyzed trials in the Eurêka and no Eurêka conditions (initial time window: $F(1,80)=0.60$, $p=0.44$; response time window: $F(1,80)=0.04$, $p=0.84$). We did not observe a significant interaction effect between the *semantic remoteness* (close or distant) and the way the subject solved the problem (with or without Eurêka) on the average number of included trials in EEG analyses, for either the initial time window ($F(1,80)=0.67$, $p=0.42$) or the response time window ($F(1,81)=0.12$, $p=0.73$) (**Figure S2**). This indicated that there is no unbalanced distribution of analyzed trials between *semantic remoteness* and *insight solving* conditions.

**Supplementary figures**

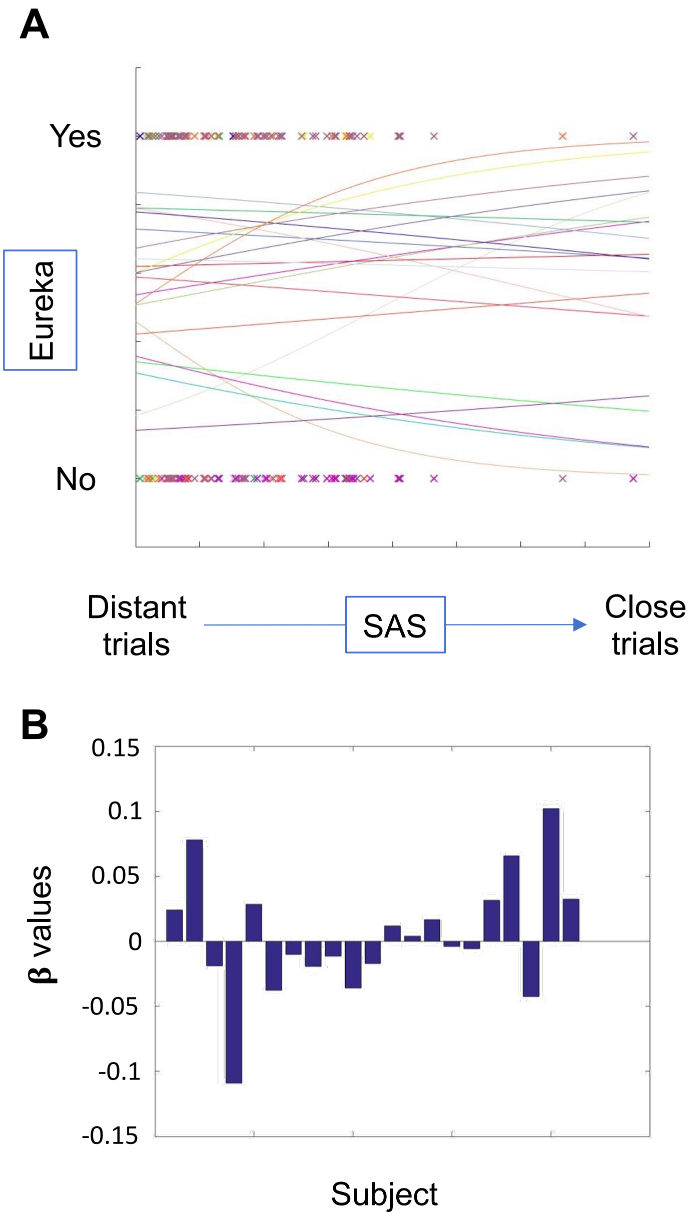


**Figure S1. A. Individual logistic regressions exploring *SAS* as a predictor of the Eurêka report.** Each cross represents a trial with its value of *SAS* (in x-axis) and the report or not of a Eurêka. (binary variable, y-axis). Each colour represents a participant (n=21) with his/her fitting curve based on individual logistic regression. None of these models was significant at the individual level (all p>0.05). **B. Bar plot of regression coefficient values.** Each bar represents a subject (n=21).

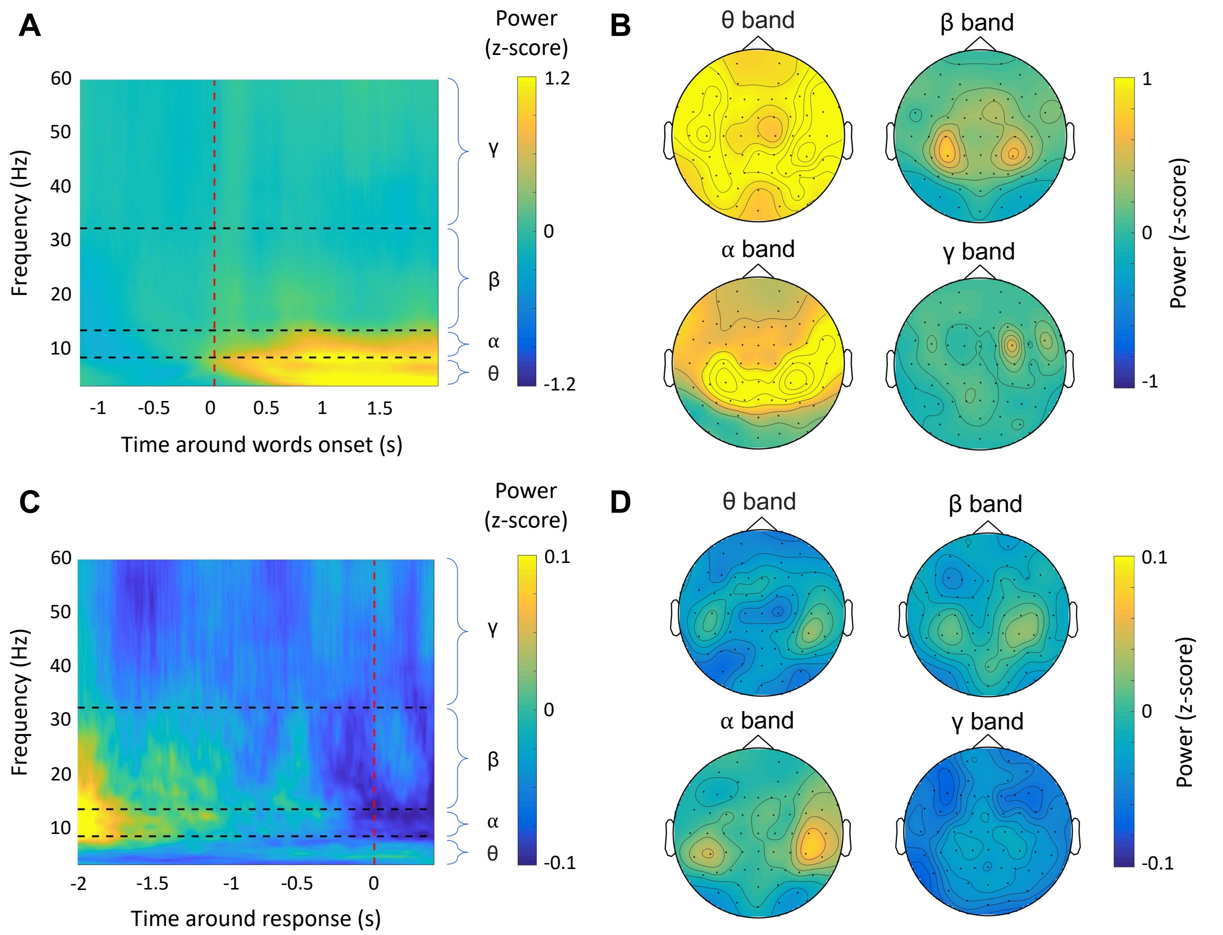


**Figure S2. Mean time-frequency maps of EEG activity across all trials. A.** Time-frequency map across all trials averaged across all electrodes and participants during the initial time window. The time course (x-axis, in seconds) was locked on the onset of the display of the cue words (time 0, vertical dashed red line). The period before time 0 corresponds to the baseline. The color bar represents the power scale (z-score of dB). Frequencies (y-axis, in Hz) were separated into four predefined frequency bands (horizontal dashed black lines): θ (3-7Hz), α (8-12Hz), β (12-30Hz), γ (31-60Hz). **B.** Topographical maps of the theta (θ, top left), alpha (α, bottom left), beta (β, top right) and gamma (γ, bottom right) bands during the initial time window (power in each band was averaged across time, from 0 to 2s). The color bar represents the power scale (z-score of dB). **C.** Same as **A** but for the response time window. The time course was locked on the participants' responses (time 0, vertical dashed red line). **D**. Same as **B** but for the response time window (power in each band was averaged across time, from -2 to 0s).

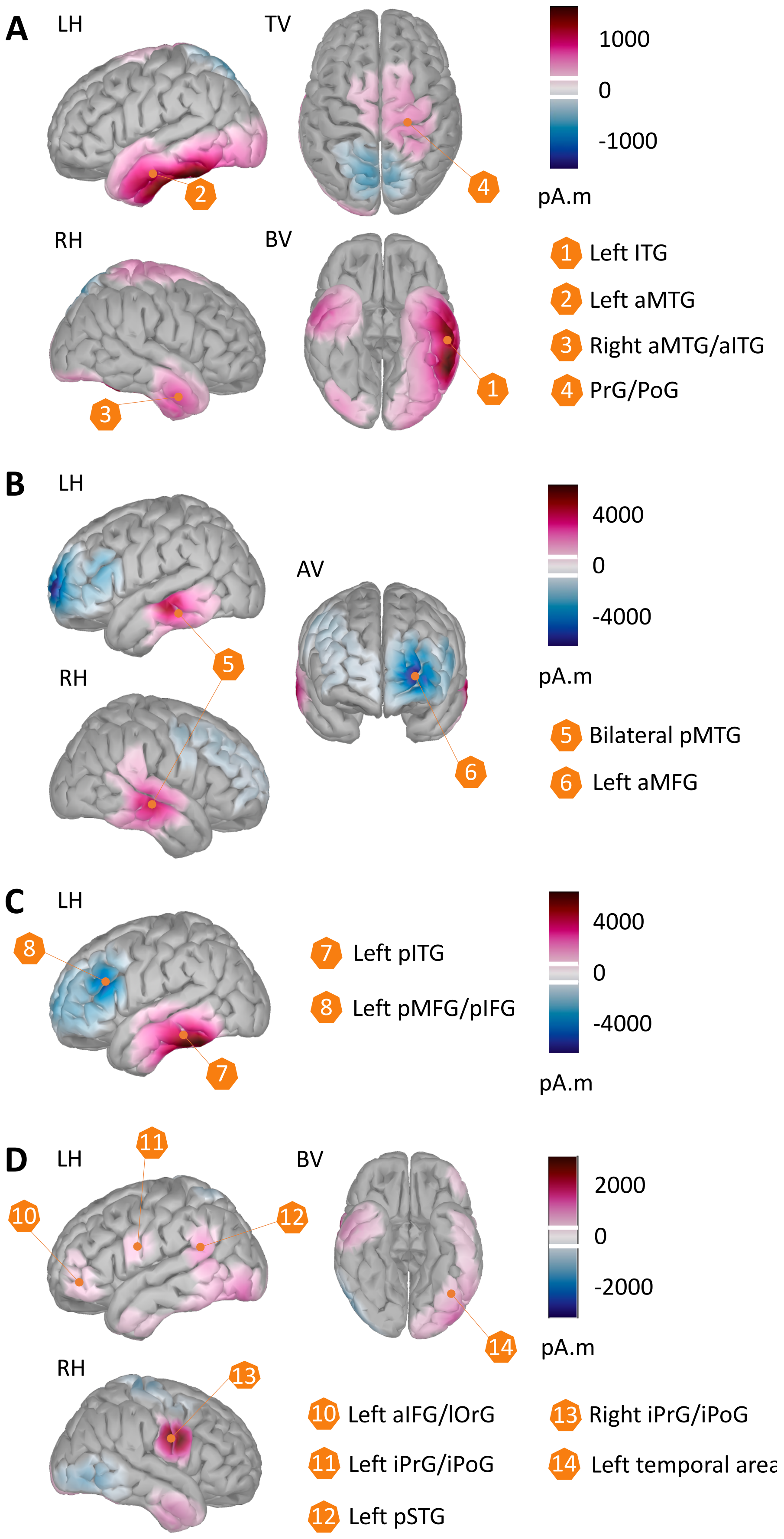
A
LH
TV
1000
0
-1000
pA.m
RH
BV
1 Left ITG
2 Left aMTG
3 Right aMTG/aITG
4 PrG/PoG
B
LH
AV
4000
0
-4000
pA.m
RH
5 Bilateral pMTG
6 Left aMFG
C
LH
7 Left pITG
8 Left pMFG/pIFG
4000
0
-4000
pA.m
D
LH
BV
2000
0
-2000
pA.m
RH
10 Left aIFG/lOrG
11 Left iPrG/iPoG
12 Left pSTG
13 Right iPrG/iPoG
14 Left temporal area

**Figure S3. Source reconstruction of the time course of EEG power related to the significant clusters obtained at the sensor level for the effect of the *remoteness* of semantic associations. A.** Source reconstruction was computed trial by trial for each individual and averaged across close and distant trials (based on the median value). Sources were then averaged across the alpha cluster's time window and frequency band found at the sensor level and across individuals. The contrast of source reconstruction (distant – close) was represented at the cortical surface of the 3D template MRI normalized in the MNI space. Values of the current density map are indicated with the color bar (in pA.m) from negative value (in blue) to positive value (in purple). The white lines in the color bar indicate the threshold used to visualize the source on the normalized brain rendering. **B.** Same as **A** for the first component of the beta cluster. **C.** Same as **A** for the second component of the beta cluster. **D.** Same as **A** for the theta cluster.

AV – Anterior view, BV – Bottom view, LH – Left hemisphere, RH – Right hemisphere, TV – Top view.

a – Anterior part, i – Inferior part, p – Posterior part.

MFG – Middle frontal gyrus, MTG – Middle temporal gyrus, IFG – Inferior frontal gyrus, ITG – Inferior temporal gyrus, OrG – Orbitofrontal gyrus, PoG – Postcentral gyrus, PrG – Precentral gyrus, STG – Superior temporal gyrus.

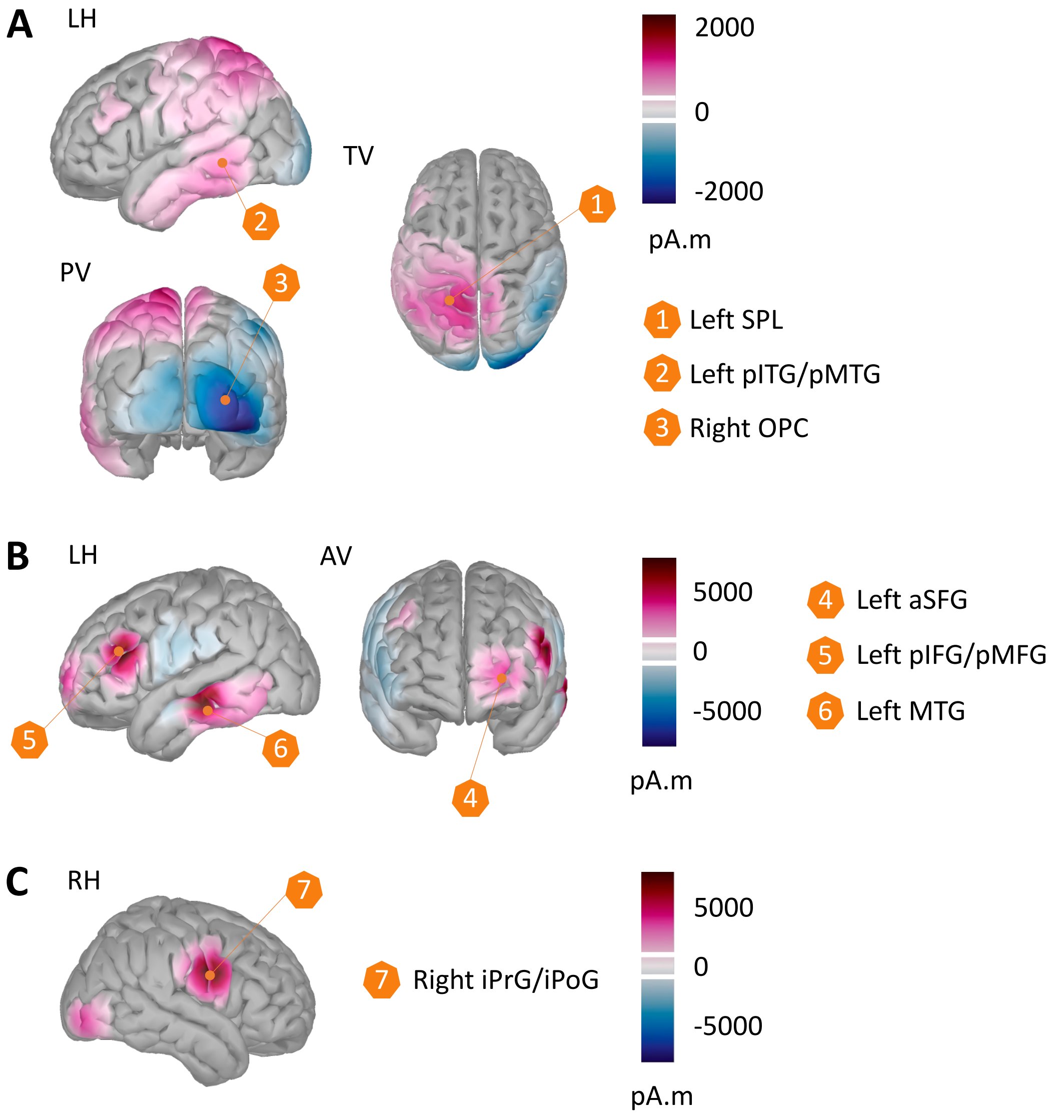


**Figure S4. Source reconstruction of the time course of EEG power related to the significant clusters obtained at the sensor level for *insight solving*. A.** Source reconstruction was computed trial by trial for each individual and averaged across trials with Eurêka and those without Eurêka. Sources were then averaged across the alpha cluster's time window and frequency band found at the sensor level and then across individuals. The contrast of source reconstruction (Eurêka – no Eurêka) is represented at the cortical surface of the 3D template MRI normalized in the MNI space. Values of the current density map are indicated with the color bar (in pA.m) from negative value (in blue) to positive value (in purple). The white lines in the color bars indicate the threshold used to visualize the source on the normalized brain rendering. **B.** Same as **A** for the gamma cluster. **C.** Same as **A** for the theta cluster.
AV – Anterior view, LH – Left hemisphere, PV – Posterior view, RH – Right hemisphere, TV – Top view.
a – Anterior part, i – Inferior part, p – Posterior part.
MFG – Middle frontal gyrus, MTG – Middle temporal gyrus, IFG – Inferior frontal gyrus, ITG – Inferior temporal gyrus, OPC – Occipital polar cortex, PoG – Postcentral gyrus, PrG – Precentral gyrus, SFG – Superior frontal gyrus, SPL – Superior parietal lobule.

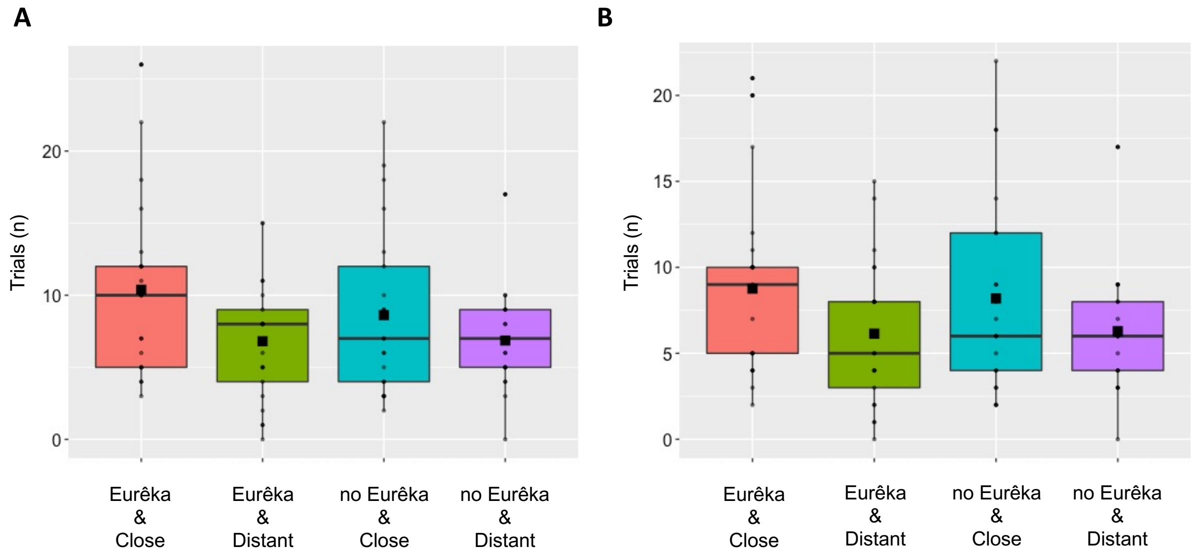


**Figure S2. Distribution of the analyzed trials across conditions.** Each box plot represents the number of trials included in the EEG analyses for close trials with Eurêka (in red), distant trials with Eurêka (in green), close trials without Eurêka (in blue), and distant trials without Eurêka (in purple). Trials were categorized as close or distant conditions depending on the median value of the *SAS*. Each point represents a participant. Color boxes represent the upper and lower quartiles. The black horizontal line represents the median value. The black filled square represents the mean of the number of analyzed trials across participants. **A** and **B** are related to the initial time window and the response time window, respectively.

**Supplementary table**

| Time window | Condition | | | |
|---|---|---|---|---|
| | Close | Distant | Eurêka | no Eurêka |
| Initial | 18.8 (6.8) | 13.3 (5.9) | 16.7 (9.3) | 15.3 (8.3) |
| | 32.1 (11.5) | | 32.0 (11.7) | |
| Response | 17.4 (6.7) | 12.9 (6.5) | 14.9 (8.5) | 14.5 (8.2) |
| | 30.3 (12.0) | | 29.4 (11.0) | |

**Table S1. Distribution of trials used in EEG analyses.** Average number (SD) of correct trials included in EEG analyses for each condition and time window after preprocessing.
